# Acrean fish diversity back in time: Paleoichthyology of the Rio Acre fossil fauna in Brazil

**DOI:** 10.64898/2026.09.27.754785

**Authors:** Gustavo A. Ballen, Karina Alencar, Fellipe Muniz, Edson Guilherme, Leonardo Aldrin, Silvia Lomba, Lucas R. Dias-Costa, Annie S. Hsiou

## Abstract

Freshwater fishes are an important component of the Neogene fossil record of South America. They have been recorded in the Acre fossil fauna, with considerable taxonomic and ecological diversity. Novel findings from the Patos locality in Acre State have produced several specimens representing eleven taxa, including the first occurrences in the fossil record of the genera *Agamyxis*, *Glanidium*, and *Trachycorystes*. These taxa comprise two orders, Characiformes and Siluriformes, with four and seven taxa respectively. The taxa found in the Patos locality are representative of typical modern Amazonian assemblages in the region’s drainages, suggesting substantial ecological conservatism. A clustering reanalysis of the updated list of the Rio Acre fossil fish fauna is consistent with previous findings in the literature regarding clustering structure in fossil assemblages during the Neogene of South America.

## Introduction

South America harbours several of the world’s biodiversity hotspots (Myers et al., 2000). Since the breakup of Gondwana, it has been essentially a continental island, with millions of years of evolutionary processes and a distinctive biota (Jaramillo, 2023). Throughout its geological history, several key processes have affected the evolution of its biological communities (Antonelli et al., 2018). These include important paleodrainage reconfigurations across the continent (Cassemiro et al., 2023), mountain uplift associated with the Andean orogeny (Gregory-Wodzicki, 2000), marine transgression-regression cycles and the development of massive lacustrine environments associated with the Pebas system (Hoorn et al., 2010), and the eventual contact with North America through the closure of the Isthmus of Panama (Bacon et al., 2015).

The continent is renowned for the remarkable diversity of several vertebrate groups, including birds (Haffer, 1990), amphibians (Vasconcelos et al., 2019), squamates (Vitt et al., 2003), and marsupials (Jansa et al., 2014). Freshwater fishes are an exceptionally diverse component of the continent’s vertebrate fauna, encompassing more than 6200 species (Albert et al., 2020; Tagliacollo et al., 2024). They occur from brackish water to nearly 4000 m above sea level and inhabit most of the continent, absent only from the dry portions of the Patagonian Peninsula. Numerous groups show extreme ecological strategies, inhabiting underground water and leaf litter and even breathing atmospheric oxygen and moving through land for some distance. However, most freshwater fishes are restricted to continental waters, with very limited dispersal potential across water divides (Albert et al., 2020). This makes them a key group for addressing biogeography questions, as they are strongly restricted to the drainages they inhabit and reflect vicariance, geodispersal, and headwater capture in their interrelationships (Albert & Reis, 2011).

Despite its enormous modern diversity, our understanding of the past diversity of many freshwater fish groups is very limited. This may partly reflect the delicate skeletons of many groups (e.g., Acestrorhamphidae), their occurrence in high-erosion environments (e.g., Astroblepidae), and the low local abundance of some taxa despite their broad geographic distributions (e.g., Scoloplacidae). Together, these factors contribute to taphonomic filters that limit the number of taxa known from fossil occurrences. Although their preservation in the fossil record is problematic, they have long been documented from several localities (Lundberg et al., 2010). Among these occurrences, the Rio Acre fossil fauna preserves a snapshot of the Amazonian biodiversity through the Neogene (Cozzuol, 2006). Numerous vertebrate groups have been reported from this fossil fauna, and it has long been known to preserve fishes (Richter, 1989). Nonetheless, previous collecting efforts and ongoing fieldwork continue to recover new specimens, expanding our knowledge of the region’s vertebrate diversity. This work reports new freshwater fish specimens recovered from the Patos locality (Cozzuol, 2006) of the Rio Acre fauna, assesses their taxonomic identity, and discusses their paleoecological and paleogeographical implications, as well as their contribution to our understanding of the paleodiversity of Amazonian freshwater fishes.

### Geological setting

The fossils described herein were recovered from the Patos locality (also known as Acre 6 and LACM 4611; Frailey, 1986; Kay & Frailey, 1993), a >100-m-long riverbank exposure along the upper Acre River, ca. 38km upstream from the municipalities of Assis Brasil (Acre State, Brazil) and Iñapari (Madre de Dios Department, Peru) (10°55′55”S, 69°5′20’’W; Figure 1). The locality is extremely rich in specimens and diversity, mostly small, isolated remains. The assemblage is numerically dominated by fishes, followed by crocodylians and turtles, whereas squamates, birds, and mammals occur less frequently (e.g., Frailey, 1986; Broin et al., 1993; Bocquentin & Melo, 2006; Kay & Cozzuol, 2006; Aguilera et al., 2008; Hsiou et al., 2010; Guilherme et al., 2023; Martucci-Neto et al., 2025).

**Figure 1.**
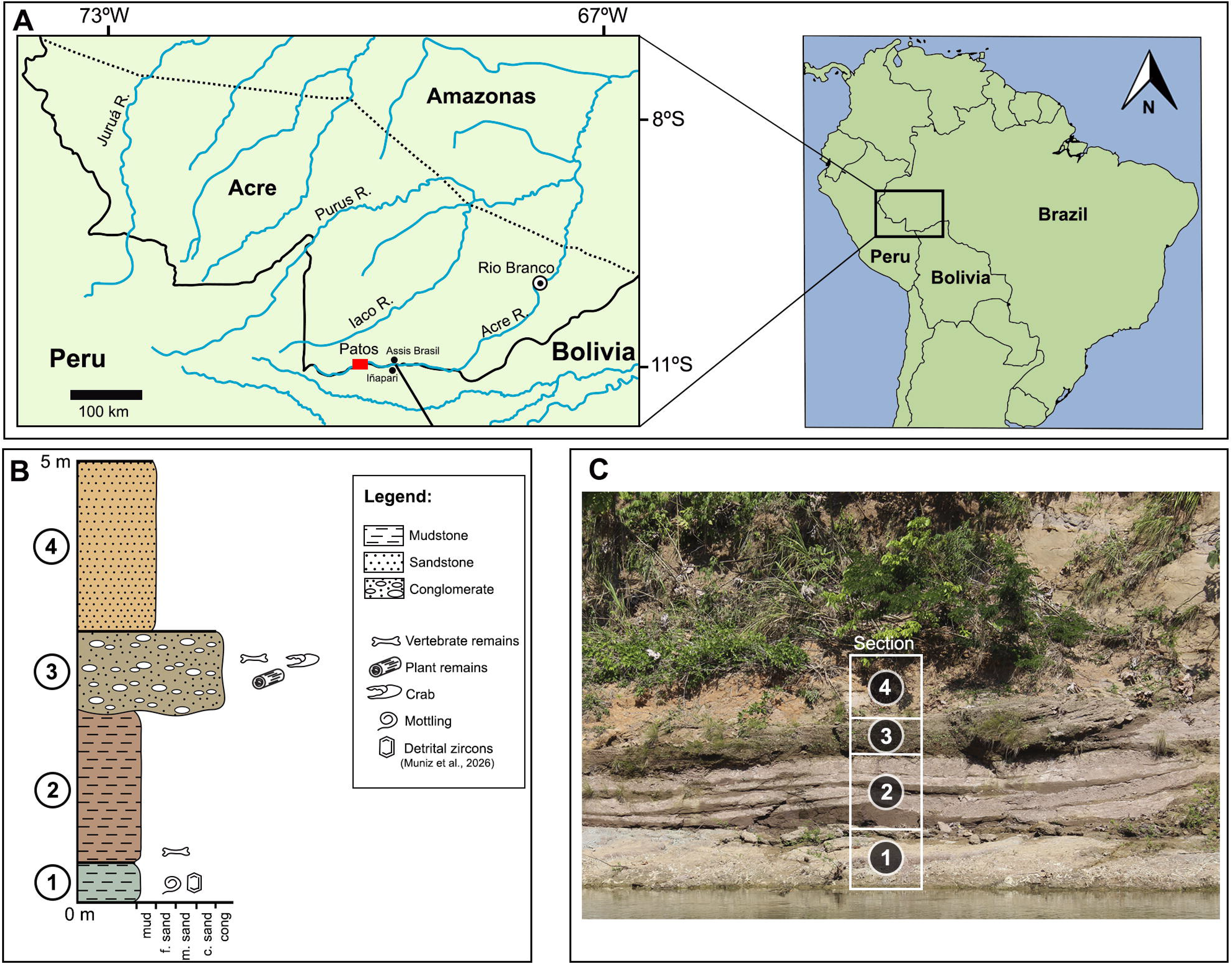
A) Map of the southwestern Amazon showing the Patos locality. B) Stratigraphic column of the outcrop exposure at the locality, part of the Solimões Formation. C) Photograph of the outgroup with the stratigraphic interpretation of the column depicted in B.

The locality’s lithology is relatively well described and resembles that of other outcrops along the Acre River (Kay & Frailey, 1993; Campbell et al., 2006; Cozzuol, 2006; Latrubesse et al., 2010). The lower portion of the outcrop, commonly submerged during the rainy season, consists of massive red- to green-colored fine-grained deposits, which are overlain by intraformational conglomerates and capped by massive to cross-bedded sandstones (Cozzuol, 2006; Latrubesse et al., 2010; Muniz et al., 2026). Nearly all vertebrate fossils, including those described herein, are concentrated within the conglomeratic deposits. The lower, fine-grained deposits contain only sparse vertebrate remains, while fossils have not yet been reported from the overlying sandstones.

The stratigraphic assignment of these fossil-bearing conglomerates remains debated (Muniz et al., 2026). Most authors regard them as part of the Solimões Formation of the Acre Basin (e.g., Bergqvist et al., 1998; Cozzuol, 2006; Latrubesse et al., 2010; Kerber et al., 2014; Guilherme et al., 2024) and interpret them as resulting from normal fluvial cut-and-fill processes (e.g., Cozzuol, 2006). Alternatively, Campbell et al. (2001, 2006) proposed that the conglomerates overlie the Ucayali Unconformity, a regional erosional surface separating the older Solimões Formation (equivalent to the Ipururo Formation in Peru) from the younger Içá Formation (equivalent to the Madre de Dios Formation in Peru). However, the existence and regional significance of the Ucayali Unconformity remain controversial (Cozzuol, 2006; Campbell et al., 2006; Muniz et al., 2026).

Despite these stratigraphic uncertainties, available biostratigraphic and geochronological evidence consistently supports a Late Miocene age for the Patos fauna. The mammalian assemblage has been assigned to the Huayquerian South American Land Mammal Age (SALMA) (Latrubesse et al., 2007, 2010), consistent with Tortonian-Messinian radiometric ages obtained from other fossiliferous outcrops of the Solimões Formation (Bissaro-Junior et al., 2019; Alvim et al., 2026). Comparable ages have also been reported from 40Ar/39Ar dating of nearby volcanic ash layers from the Madre de Dios Formation in Peru, indicating that the regional fossiliferous conglomerates are younger than ∼9 Ma (Campbell et al., 2001). In addition, the occurrence of the spore *Cyatheacidites annulatus* at Patos suggests an age not older than ca. 6 Ma based on the pollen zonation proposed for the Solimões Formation by Jaramillo et al. (2024) and the occurrence reported by Leite et al. (2021). Recent CA-TIMS U-Pb ages from detrital zircons from Patos further support this interpretation, indicating a maximum depositional age of 5.573 ± 0.019 Ma for the fossil-bearing deposits (Muniz et al., 2026).

## Materials and methods

Anatomical terminology follows Ballen et al. (2022) for Serrasalmid teeth, Ballen et al. (2021) for Cynodontid osteology, and Ballen & de Pinna (2022, 2024) for Siluriform appendicular terminology. Extant and extinct comparative material as reported in Ballen & de Pinna (2022), and Ballen et al. (2021, 2022, 2026). A slightly modified version of the dataset in Ballen et al. (2026) including the new records was subject to the same clustering analysis described in that paper, and the results are available as Supplementary material. Code and data are available both on github (https://github.com/gaballench/patos_rio_acre_fossil_fishes) and on zenodo (DOI: 10.5281/zenodo.23931068). The analysis was carried out in R v4.5.x (R Core Team, 2026) with the package pvclust v2.2 (Suzuki et al., 2019).

## Results

Actinopterygii Woodward, 1891

Characiformes Fink & Fink, 1981

Cynodontidae Eigenmann, 1907

*Hydrolycus* Müller & Troschel, 1844

*Hydrolycus* sp.

Figure 2A-B.

**Figure 2.**
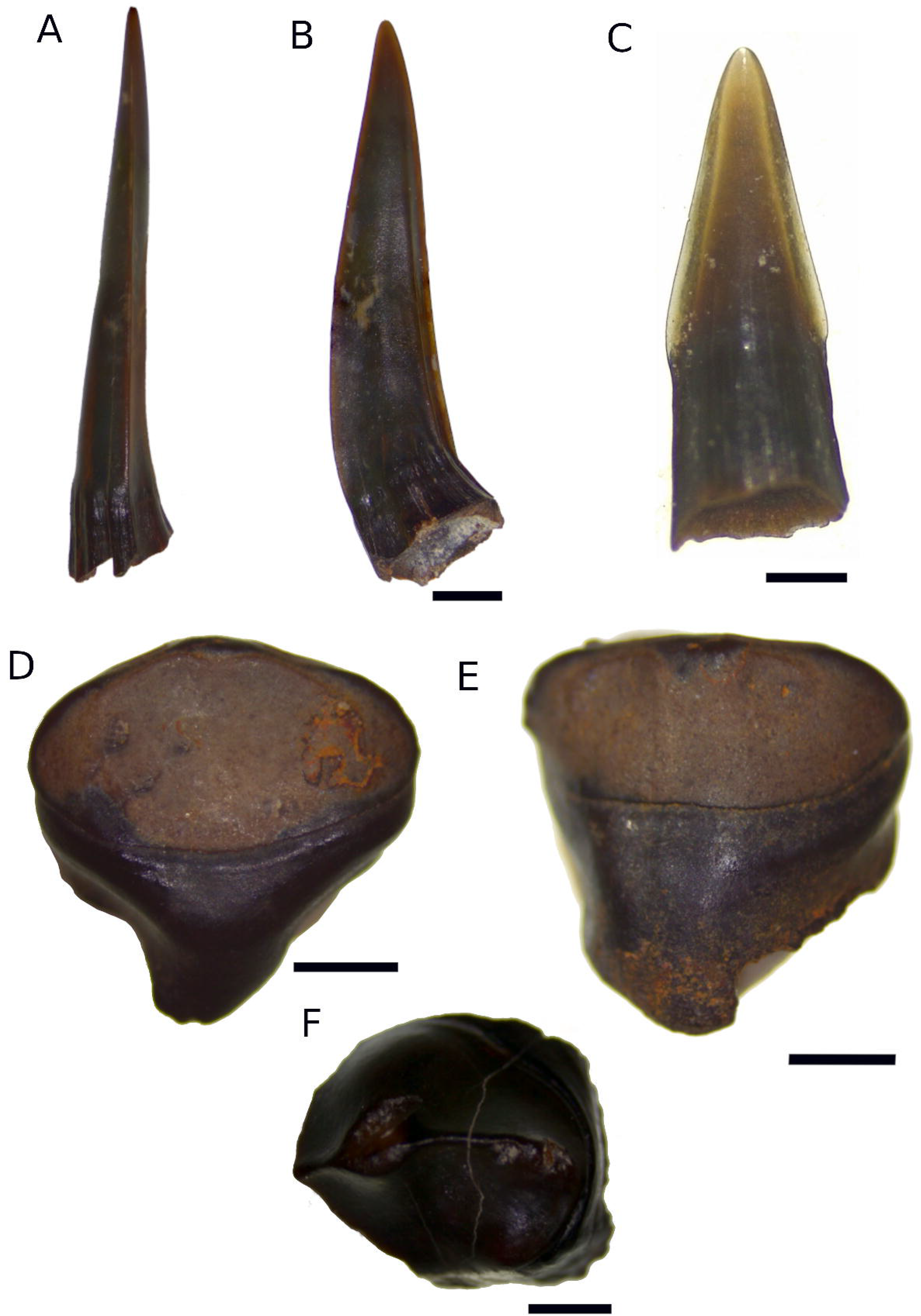
Characiformes from the Patos locality. *Hydrolycus* sp., A) leading canine in anterior view, and B) lateral view. cf. *Hoplias* sp., C) isolated tooth in lingual view. *Colossoma* sp., D) isolated tooth D4 or D5 in occlusal view, and E) in labial view. cf. *Piaractus* sp., F) isolated tooth PM5 in occlusal view. All scale bars are 2mm except in C where it represents 0.5mm.

Referred material: Leading dentary tooth, UFAC 7156.

### Osteological description

Strong dentary tooth with labio-lingual curvature. Enameloid preserved from base to tip. Insertion plane commisuro-symphyseal. Three cutting edges, two on margins of lingual surface and one on labial surface. Preserved tooth base with vertical fine ridges.

### Remarks

Due to the insertion plane of the tooth, possible species are *Hydrolycus aramatus*, *H. scomberoides*, *H. tatauaia*, or yet another undescribed species. *Hydrolycus wallacei* differs from these in that it presents a labiolingual insertion plane (Ballen et al., 2021). All species are restricted to drainages east of the Andes, and one extant species, *Hydrolycus scomberoides*, is known from the Pliocene of northern Colombia (Ballen et al., 2021).

Erythrinidae Valenciennes in Cuvier & Valenciennes, 1847

*Hoplias* Gill, 1903

cf. *Hoplias* sp.

Figure 2C.

Referred material: Isolated tooth preserving all of the enameloid and part of the root, UFAC 8618.

### Osteological description

Lanceolate tooth with cutting edges on commisuro-symphyseal edges. Enameloid extending onto tooth base on both symphyseal and commisural margins. Overall, enameloid surface smooth.

### Remarks

The fossil record of *Hoplias* is mostly Neogene in South America and is well known from the Acre fauna (Lundberg et al., 2010). The genus includes a modest number of extant species. It seems to show geographic structuring, although high variability in morphology and chromosome number obscures possible structuring across species, at least within the *Hoplias malabaricus* complex (Cardoso et al., 2018).

Serrasalmidae Bleeker, 1859

Colossomatinae Kolmann, Hughes, Hernandez, Arcila, Betancur-R, Sabaj, López-Fernández & Ortí 2021

*Colossoma* Eigenmann & Kennedy, 1903

*Colossoma* sp.

Figure 2F.

Referred material: Isolated tooth D4 or D5, UFAC 8619.

### Osteological description

Globose isolated tooth with occlusal surface deflected lingually— cutting edge raised and continuous from commisural to symphyseal margins.. Occlusal surface somewhat concave distally and then convex basally, more pronounced than labial surface, with small concavity on lingual margin of tooth base.

### Remarks

Serrasalmid isolated teeth have been recorded from the Paleocene of Bolivia to the Pliocene of Colombia (Ballen et al., 2026). They are a common component of vertebrate microremains in South America due to the robustness of their teeth. Ballen et al. (2022) provided several diagnostic characters. They discussed the potential of dental characters for identifying multiple occurrences of the family in the fossil record, showing how serial homology and heterodonty are useful for this task. However, a thorough examination of this anatomical complex is still needed to reassess the many specimens already reported from across South America.

*Piaractus* Eigenmann, 1903

cf. *Piaractus* sp.

Figure 2D-E.

Referred material: Isolated tooth PM5, UFAC 8620.

### Osteological description

Premaxillary tooth PM5 with compressed base and robust crown. Occlusal surface roughly oval in outline, with worn cutting edge. Apparently unicuspid. Labial surface with middle vertical elevated area and two lateral slightly concave areas. Enameloid wider than high in labial view. Tooth base with lateral constrictions, both in labial and occlusal views.

### Remarks

The specimen mostly agrees with the characters provided by Ballen et al. (2022) for the genus *Piaractus*. However, an in-depth review of comparative anatomy is still needed to provide a more precise identification for this specimen. The genus comprises three species distributed in drainages east of the Andes: *P. brachypomus* in the Amazon drainage, P*. mesopotamicus* in the Paraná-Paraguay drainage, and *P. orinoquensis* in the Orinoco drainage (Escobar et al., 2019).

Siluriformes Fink & Fink, 1981

Auchenipteridae Bleeker, 1862

*Glanidium* Lütken, 1874

cf. *Glanidium* sp.

Figure 3A-B.

**Figure 3.**
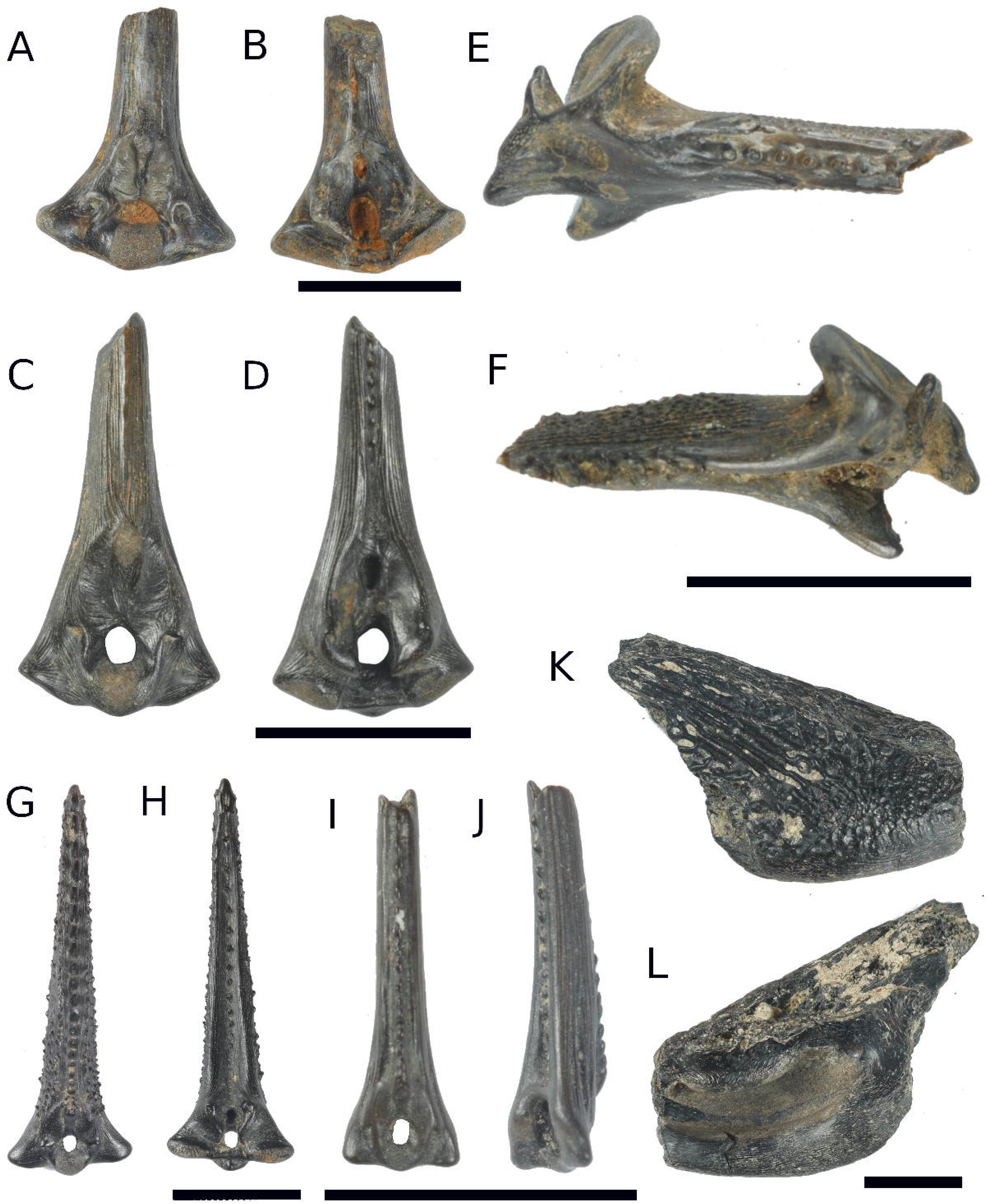
Siluriformes except *Phractocephalus* from the Patos locality. cf. *Glanidium* sp., A) dorsal-fin spine in anterior view and B) posterior view. *Lophiosilurus* sp., C) dorsal-fin spine in anterior view and D) posterior view. *Trachycorystes* sp., E) pectoral-fin spine in anterior view and F) posterodorsal view. *Agamyxis* sp., G) dorsal-fin spine in anterior view and H) posterior view. *Hemidoras* sp., I) dorsal-fin spine in anterior view and J) posterior view. cf. *Oxydoras* sp., K) partial cleithrum in lateral view and L) mesial view. All scale bars are 10mm.

Referred material: Dorsal-fin spine base, UFAC 8621.

### Osteological description

Spine base wider than high, triangular in outline. The anterior articular surface finely rugose, roughly square in outline. Triangular foramen in anterior view,base width twice its height. Anterior fossae lateral to foramen, circular in outline, and with diameter about the same as foramen’s height; elevated bony ridge on dorsomedian margin of fossa. Attachment surface for *ligamentum interspinalis is* high and oval in outline, around three times as high as wide, spanning from dorsal margin of foramen to inflection point of the anterior longitudinal ridge. Lateral fossae present, thin and oval in outline. Lateral condyle short, with ventral and dorsolateral margins straight, projected laterally; posterior expansion modest. Lateral articular surfaces low and oval in outline; lower medially, higher laterally, and finely granular in texture. Posterior process short and wide, oblique and ridge-like, with lowest point closest to midline, uppermost point closest to lateral margin of spine base. Posterior fossae thin and oblique, following orientation of posterior process. Posterior sulcus surrounded by flat-surfaced hemitrichia. Anterior and posterior ornament absent.

### Remarks

Dorsal-fin spines with very wide bases are typical of the Auchenipteridae, and among these, the genus *Glanidium* best fits the characters observed in this specimen. However, their taxonomic limits within the Auchenipteridae remain to be characterized, and therefore, we prefer to assign them to that genus tentatively. The extant species of *Glanidium* are present in drainages peripheral to the Amazon-Orinoco area, including coastal rivers of the Guiana Shield, the São Francisco River basin, the Paraná River basin, and coastal drainages of southern Brazil from the states of Espírito Santo to Rio Grande do Sul (Sarmento-Soares & Martins-Pinheiro, 2013). If the specimen herein studied in fact represents a species of *Glanidium*, it would be the first record in the Amazon Basin, suggesting a historically wider geographic distribution and potentially the disjoint distributions of the extant species.

*Trachycorystes* Bleeker, 1858a

*Trachycorystes* sp.

Figure 3E-F.

Referred material: Pectoral-spine base, UFAC 8623.

### Osteological description

Left pectoral-fin spine preserving most of base and less than proximal 1/3 of spine shaft. Median process well developed, acute, and continuous with dorsal surface of articular facet for scapulocoracoid. Anterior process present, with anterior emargination in anterior view and a spongy anterior surface. Anterior fossa oval with horizontal axis largest. Flange of anterior process absent. Anteroventral emargination wide, not strongly concave, with ventral margin slightly straight in anterior view. Basal recess oval almost completely covered by sediment in posterior view. Posterior process nearly absent, posterior keel smooth and not very prominent. Dorsal process well developed, deflected laterally in anterior view, discoid in mesial view. Anterior shaft process consists of tubercles basally, then apparently antrorse spinules, but their tips are not preserved. Posterior ornament consisting of retrorse blades, partly fractured, but with vertical axis largest in cross-section. Dorsal and ventral shaft surfaces with fine ridges, covered densely with acute tubercles, more densely on the proximal region. Basal-most portion of shaft expanded on antero-posterior axis.

### Remarks

The finely acute tubercles covering the dorsal and ventral surfaces of the spine shaft, and the well-developed median process, directed dorsally and pointed, are characteristics of the genus *Trachycorystes* amongst Neotropical Siluriformes. Spines in the genus *Trachelyopterichthys* have a series of basally fused anterior straight spinules (Ballen et al., 2026), which are different from the individual spinules seen in *Trachycorystes*. The spine base is less robust in the genus *Trachelyopterus*, without the basal anteroposterior expansion seen in *Trachycorystes*. This is the first fossil record of the genus, which is endemic to the Neotropics and widely distributed in the Amazon and Orinoco drainages (Britski & Akama, 2011).

Doradidae Bleeker, 1858b

*Agamyxis* Cope, 1878

*Agamyxis* sp.

Figure 3G-H.

Referred material: Dorsal-fin spine, UFAC 8624.

### Osteological description

Complete dorsal-fin spine. Anterior articular surface finely rugose, nearly round in outline, with dorsal emargination in anterior view. Foramen subovoid in anterior view, with width just less than height. Anterior fossae located lateral to basal half of foramen, round in outline, deflected ventrally with bony rim of pointy appearance, and with diameter of about 1/4 of foramen height. Attachment surface for *ligamentum interspinalis* high and teardrop in outline, around three times as high as wide, spanning from dorsal margin of foramen to dorsal margin of inflection point of anterior longitudinal ridge. Lateral fossae absent. Lateral condyle with moderate lateral projection, with ventral and dorsolateral margins straight; posterior expansion very short. Lateral articular surfaces are low and oval in outline, with finely granular texture. Posterior process short and with ridge from upper lateral portion to lower median. Posterior fossae wide, following posterior process outline. Dorsal margin of lateral articular surfaces highest laterally, lowest medially. Posterior sulcus very wide, with concave-surfaced hemitrichia and surrounded by sharp longitudinal keel on each hemitrichium, forming very wide sulcus concave on the hemitrichia. Shaft preserving anterior, posterior, and lateral ornaments, as well as fenestrae communicating interior of the shaft lumen with epidermal tissue. Anterior ornament consisting of multicuspidate complex on inflection point of anterior longitudinal ridge and three to four subsequent ornaments dorsally, then becoming curved antrorse spinules throughout length of spine; ornaments becoming larger distally; spinules closely set proximally, leaving space inbetween less than spinule diameter, and then becoming more spaced, with space for two or more spinules inbetween distalmost ornaments. Posterior ornament absent on proximal 1/5 of spine length, then consisting of sharp tubercles, with space in between for three to five ornaments. Lateral ornament consisting of subparallel ridges and numerous small spinules, wider than surface ridges and also oriented obliquely, similar to antrorse ornaments, then becoming tubercles distally perpendicular to spine surface plane distally. Shaft outline triangular in transversal view.

### Remarks

This is the first known fossil record for the genus *Agamyxis*. The two known extant species are found in the Amazon and Orinoco drainages (Birindelli, 2014). The combination of spinules on the lateral surfaces of the spine shaft, antrorse spinules on the anterior surface, and tubercles on the posterior surface separates *Agamyxis* from other Doradids, in particular from *Acanthodoras* (ornament on posterior surface absent), and *Amblydoras* (ornament on both anterior and posterior surfaces absent). Ballen & de Pinna (2022, p. 619) offered an overview of the presence of spinules or tubercles on shaft surfaces, especially in the Auchenipteridae, Ariidae, and Doradidae.

*Hemidoras* Bleeker, 1858a

*Hemidoras* sp.

Figure 3I-J.

Referred material: Dorsal-spine, UFAC 8622

### Osteological description

Incomplete dorsal-fin spine, about half shaft preserved. Base complete. Anterior articular surface finely rugose, roughly square in outline. Foramen triangular in anterior view, with width half height. Anterior fossae located lateral to basal half of foramen, oval in outline, and with diameter about half of foramen height; elevated bony ridge on the dorsomedian margin of fossa, with triangular appearance. Attachment surface for *ligamentum interspinalis* high and oval in outline, around three times as high as wide, spanning from dorsal margin of foramen to inflection point of anterior longitudinal ridge. Lateral fossae present, thin and oval in outline. Lateral condyle short and ventrally deflected, with ventral and dorsolateral margins straight; posterior expansion very short. Lateral articular surfaces are low and subtriangular in outline, with a finely granular texture. Posterior process very short and eroded. Posterior fossae poorly preserved due overall erosion of specimen. Posterior sulcus surrounded by round-surfaced hemitrichia. Shaft preserving anterior, posterior, and lateral ornaments, as well as fenestrae communicating interior of shaft lumen with epidermal tissue. Anterior ornament consisting of antrorse spinules from inflection point of anterior longitudinal ridge up to preserved dorsalmost portion of shaft; spinules closely set, leaving space inbetween less than spinule diameter. Posterior ornament absent on proximal 1/4 of preserved spine length, then consisting of antrorse spinules, apparently flat, with space inbetween measuring equal to spinule diameter or more. Lateral ornament consisting of subparallel ridges. The shaft outline quadrangular in transversal view.

### Remarks

The specimen seems to have been transported, as indicated by the significant surface erosion, particularly at the base of the spine. It is the second fossil record of the genus from Neogene strata in South America. The genus was first reported from the Pliocene Ware Formation in northern Colombia (Ballen et al., 2026) and is now also present in the Acre fauna. Ballen et al. (2026) discussed the interrelationships of *Hemidoras*. They provided a set of diagnostic characters for the genus amongst the Doradidae, which are present in this specimen from the Acre fauna. The genus *Hemidoras* comprises five species, all distributed in the Amazon and Orinoco drainages (Birindelli, 2014).

*Oxydoras* Kner, 1855

cf. *Oxydoras* sp.

Figure 3K-L.

Material examined: Partial cleithrum, UFAC 7215.

### Osteological description

Right cleithrum preserving lateral swollen region of insertion for pectoral-fin spine and incomplete part of posterior process. Ornament consisting of radiating irregular ridges and tubercles with focus on lateralmost point of the swollen region.

### Remarks

The lateral ornament of the cleithrum is well developed and consisting of ridges and tubercles in *Oxydoras*. The condition present in our specimen is similar to that genus but it is necessary to compare with other genera in order to verify the generic identity of the specimen. The genus *Oxydoras* is widely distributed in the Amazon and Orinoco drainages and has been previously reported from Neogene strata of South America (Lundberg et al., 2010; Ballen et al., 2026). Representatives of *Oxydoras* tend to inhabit dusky waters, such as floodplain lakes, and can reach large sizes, exceeding 1 m in standard length (Sabaj & Arce-H, 2021).

Pimelodidae Bonaparte, 1835

*Phractocephalus* Agassiz in Spix & Agassiz, 1829

*Phractocephalus acreornatus* Aguilera, Bocquentin, Branco, Lundberg & Maciente, 2008

Figure 4.

**Figure 4.**
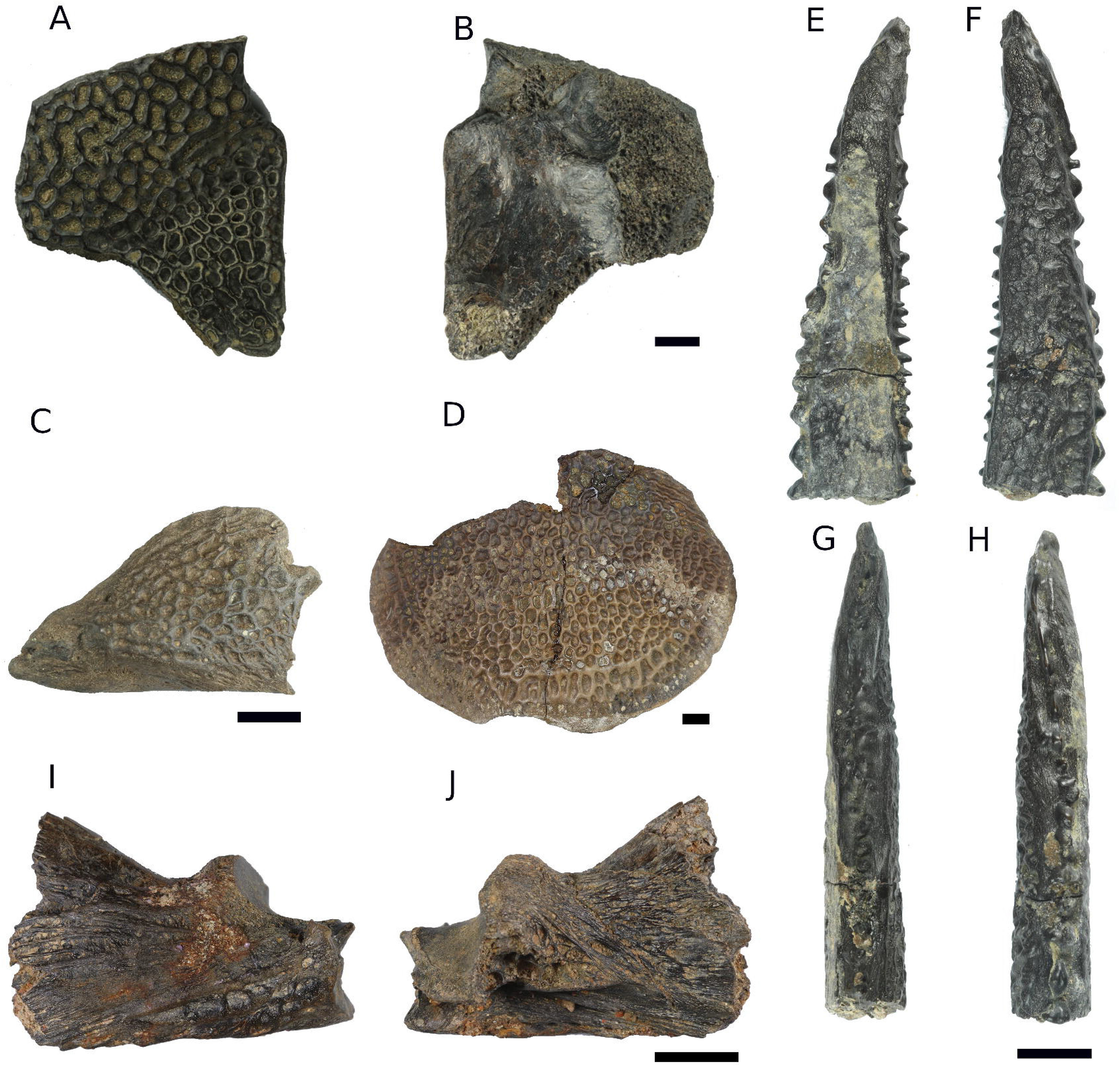
*Phractocephalus acreornatus* from the Patos locality. Pterotic bone in A) dorsal view and B) ventral view. Interopercle C) in lateral view. Posterior process of supraoccipital bone in D) dorsal view, anterior upwards. Pectoral-fin spine shaft in E) ventral, F) dorsal, G) posterior, and H) anterior views. Left anguloarticular in I) lateral view and J) Mesial view. All scale bars are 10mm.

Referred material: Right pterotic fragment, UFAC 7162. Left interopercle, UFAC 7058. Left pectoral-fin spine fragment, UFAC 7068. Posterior process of supraoccipital, UFAC 6997.

### Osteological description

Pterotic preserves most of dorsal surface, from anterior concave margin at level of articular facet for hyomandibula to posterior level of emargination for articulation with posttemporo-supracleithrum. Articulations with sphenotic, extrascapular, and supraoccipital bones preserved dorsally. The ventral surface relatively well preserved, with spongy bone exposed along articulation with exoccipital (not preserved). Dorsal surface of pterotic richly ornamented with reticulating ridges and few tubercles. Interopercle preserves lateral surface but not margins or mesial surface, which shows exposed spongy bone. Bone surface consists of fine to coarse reticulating ridges on posterior 3/4, remaining surface being smooth and apparently covered by connective tissue and skin. Posterior process of supraoccipital ovoid in outline, with lateral and posterior margins well preserved. Anterior portion continuous with body of supraoccipital missing. Anterolateral surface with oblique well-defined ridges, remainder of surface strongly reticulate with pits of oval, round, and polygonal outline. Surface ornament less developed on marginal portion of process but still visible. General surface of posterior process of supraoccipital widely convex but without evident medial keel. Left anguloarticular preserving articular facet for quadrate posteriorly, to beginning of ascending process where dentary articulates with anguloarticular anteriorly. Anterior lateral surface ornamented with parallel ridges; posterior lateral surface ornamented with parallel ridges showing reticulations and oval to circular cavities at level of articular facet for quadrate. Articular facet for quadrate bilobed in outline, with ventral longitudinal rim. Posterior projection of anguloarticular past articular facet for quadrate absent. Posterior process of anguloarticular present and straight. Pectoral-fin spine preserving distal 1/4 or 1/5, well-preserved, tip intact. Ornament present on anterior, posterior, dorsal, and ventral surfaces. Distal ornament highly irregular at growth zone. Dorsal ornament more developed than ventral one. Anterior ornament consists of blunt, wide tubercles. Posterior ornament consists of sharp spinules, from uniserial to distally irregular biserial, sometimes with tip deflected retrorsely. Because of the preserved length of about 6.5cm, otal spine length should have been around 30cm.

### Remarks

*Phractocephalus acreornatus* was described from the Acre fauna in Brazil (Aguilera et al., 2008). Our specimens match the original description, and the ornament on the dorsal surface of the posterior process of the supraoccipital, showing oblique straight ridges, matches the condition seen in *P. acreornatus* (cf. fig. 7 in Aguilera et al., 2024), despite other diagnostic characters not preserved in our specimens. Therefore, we assign it to this species rather than to other fossil congeners, such as *P. yaguaron* and *P. nassi* (Ballen et al., 2026), or to the extant *P. hemioliopterus*. However, the interopercle herein described differs in shape from the specimen described in Aguilera et al. (2008), which may be due to size differences, as the interopercle in the latter reference is larger than ours. The spine herein described differs from the paratype in Aguilera et al. (2008, fig. 10) in that their specimen is considerably flatter, which may be due to larger size or to the preserved portion being more distal in ours, thus precluding estimation of the true spine width.

Pseudopimelodidae Fernández-Yépez & Antón, 1966

*Lophiosilurus* Steindachner, 1876 *sensu* Shibatta et al. (2021)

*Lophiosilurus* sp.

Figure 3C-D.

Material examined: Dorsal-fin spine, UFAC 8625.

### Osteological description

Dorsal-fin spine base and proximal part of spine shaft preserved. Spine base width 2/3 of base height; outline triangular. Anterior articular surface finely rugose, oval in outline, and with dorsal emargination in anterior view. Foramen oval in anterior view, with base width 2/3 height. Anterior fossae located lateral to foramen, oblique oval in outline with elevated bony ridge on dorsomedian margin, sigmoid, and with height about the same as foramen height. Attachment surface for *ligamentum interspinalis* massive, half-moon shaped, and spans from foramen to inflection point of anterior longitudinal ridge, around three times higher than wide. Lateral fossae absent. Lateral condyle very short, with straight, laterally projecting ventral and dorsolateral margins; posterior expansion very short, straight, and projected laterally. Lateral articular surfaces low and widely oval in outline; lower medially, higher laterally, and finely granular in texture. Posterior process short and wide, oblique and ridge-like, with lowest point closest to midline, uppermost point closer to lateral margin of spine base. Posterior fossae thin and oblique, following orientation of posterior process. Posterior sulcus surrounded by hemitrichia with surface ornamented by fine longitudinal ridges. Ornament on anterior longitudinal ridge absent; ornament on posterior sulcus consisting of flat, blunt tubercles, starting halfway through preserved shaft length, absent on shaft section proximal to base.

### Remarks

Recently, phylogenetic analyses synonymized the genera *Cephalosilurus* and *Lophiosilurus* (Shibatta et al., 2021). Among the five extant species, the specimen most resembles *Lophiosilurus alexandri* because of its very wide, robust dorsal-fin spine, consistent with a very depressed body, a characteristic of the species among Pimelodoid Catfishes. The extant *L. alexandri* is restricted to the São Francisco River basin in Brazil. Lundberg et al. (2010, figure 17.1) reported an isolated dorsal-fin spine (LACM 128374) of either *Cephalosilurus* or *Pseudopimelodus* from the Rio Acre fauna in Madre de Dios, Perú. Later, Shibatta et al. (2021) reidentified it as either *L. alexandri* or *L. fowleri* based on the high base-to-length ratio and general robustness, unlike other genera in the family. Our specimen is in most details identical to LACM 128374, and we consider them to be conspecific. Shibatta et al. (2010) suggest that LACM 128374 may represent an undescribed species that needs confirmation through thorough comparisons between the fossil specimens and comparative material of the genus *Lophiosilurus*.

## Discussion

### Previous fish occurrences in the Acre fauna and new records

Several fish taxa have been recovered from the Rio Acre fossil fauna. Aguilera et al. (2024) provided the most complete and up-to-date inventory of works reporting fish remains from multiple localities in Acre State from Neogene strata and list 19 taxa. However, several are identified only at the family level, and five Loricariid records come from an unpublished dissertation; therefore, we prefer to exclude these from the known Rio Acre fauna until they are formally published, reducing it to nine taxa known at least to genus level (Figure 5). Most records, however, come from Lundberg et al. (2010), which presents an overview of the fossil fishes of South America but numerous new records are also reported succinctly. Here we record eleven freshwater fish taxa for the Patos locality, three of which also appear in Aguilera et al.’s list, resulting in 17 taxa identified to genus level for the Rio Acre fossil fauna. Aguilera et al.’s (2024) record of cf. *Cephalosilurus* comes from Lundberg et al. (2010) but overlooks the subsequent reidentification of that same specimen as *Lophiosilurus* instead by Shibatta et al. (2021), which we follow herein. Records identified to family level are quite often uninformative about ecology, biogeography, and interrelationships because many freshwater fish families do show considerable variation in ecology and distribution (Albert et al., 2020; Albert & Reis, 2026).

**Figure 5.**
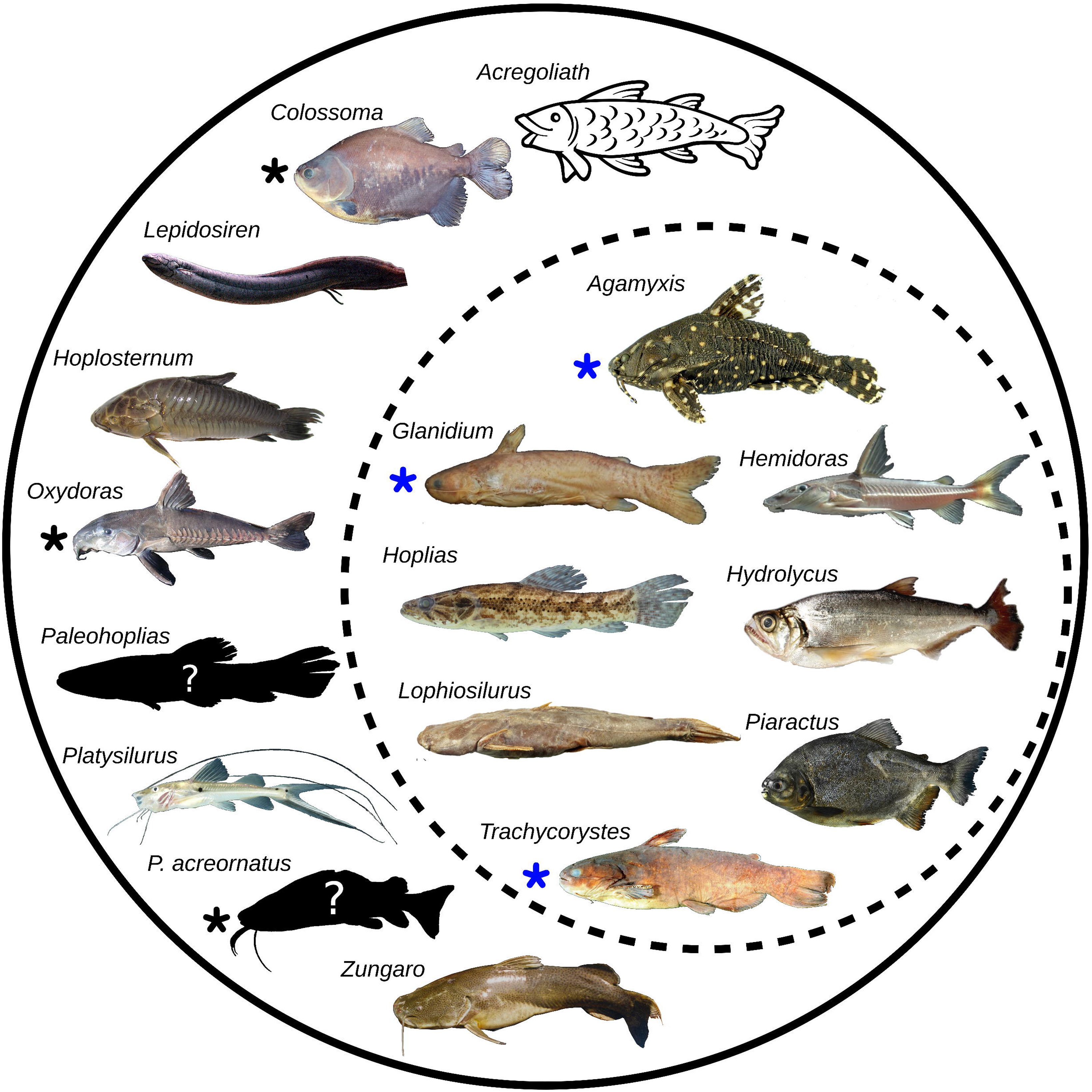
Updated fossil freshwater fish records from the Rio Acre fauna. The continuous circle represents our known boundaries in composition with 17 taxa identified at least to genus level, whereas the subset defined by the dashed circle represents the new records herein reported. Taxa identified by a black asterisk are already known from the Rio Acre fauna but also reported in this study, whereas blue asterisks within the new records are also the first known occurrences of the taxa in the fossil record. Note that *Acregoliath* is only known from scales and no certain interrelationships have been proposed for it (Richter, 1989; Ballen & Moreno-Bernal, 2019). Dark outlined fishes are fossil species for which no reconstructions are available. All the photographs are made available with appropriate licenses as detailed in the acknowledgments.

### Broader comparisons in South America

The fossil freshwater fish record in South America is long and rich. Ballen et al. (2026) summarized the known composition of Cenozoic faunas and carried out a clustering analysis to identify patterns in the fossil record. They recorded 56 freshwater taxa (including those also found in brackish and saltwater environments). Here we report four taxa not recorded in any other fossil fish fauna in South America (Figure 5). This raises the freshwater fish taxa known from the Neotropics to 60. The composition of the Rio Acre fossil fauna is typical of Amazonian assemblages; however, if our identification of cf. *Glanidium* is correct, it would represent the first Amazonian record of the genus, which is currently found in coastal Atlantic drainages of Brazil and the Guianas, as well as in the Paraná drainage (Sarmento-Soares & Martins-Pinheiro, 2013). A reanalysis of the clustering structure of fossil assemblages using the data and code in Ballen et al. (2026), adding the taxa recorded herein, resulted in essentially the same results as those of these authors: Rio Acre clusters together with the Villavieja and La Victoria formations from the Honda Group, La Venta fossil fauna in Colombia, and the Ituzaingó Formation in Argentina (Figure S1).

### Fish Paleoecology

The fish taxa found in the Patos locality suggest typical conditions of the Amazonian rainforest of South America, with the presence of large rivers or lakes, as has been interpreted in other Neogene faunas of the continent, such as the La Venta fauna (Carrillo-Briceño et al., 2023), the Castilletes Formation, and the Ware Formation (Ballen et al., 2022, 2026), and the Contamana fauna (Antoine et al., 2016). Frugivores such as *Colossoma* and *Piaractus* are typical of waterbodies with good riparian forest cover, while large catfishes such as *Phractocephalus* indicate large channels. Most other elements are either generalists or small enough to inhabit both large and small waterbodies (e.g., *Oxydoras*). Notably, most fossil occurrences come from groups found in the present composition of Amazonian basins. For instance, the fossil assemblage composition is strikingly similar to the ichthyofauna of the Sandy Beaches of the Acre River (Silva et al., 2020), showing considerable local ecological conservatism in this area of the Amazon.

## Acknowledgments

GAB was funded through FAPESP grants and fellowships (2014/11558-5, 2016/02253-1, and 2023/07838-1). GAB thanks M. Pastana for discussions on Ostariphysan osteology, especially regarding the homology of elements of the posterior portion of the cranium in Siluriformes. This research was initially supported by the Fundação de Amparo à Pesquisa do Estado de São Paulo (FAPESP; Proc. No. 2019/14153-0), awarded to ASH. We also acknowledge support from CNPq through the *Expedições Científicas – Iniciativa Amazônia +10* (Proc. No. 425126/2025-0 awarded to EG), as well as additional funding from FAPESP (Proc. No. 2025/01077-4 awarded to ASH), within the same national funding framework coordinated by CNPq and the Conselho Nacional das Fundações Estaduais de Amparo à Pesquisa (CONFAP). This research was also supported by the Conselho Nacional de Desenvolvimento Científico e Tecnológico (CNPq Process no. 406902/2022-4 INCT PaleoVert and 301579/2025) to ASH. We thank Wikicommons for providing access to fish photographs of [Colossoma](https://commons.wikimedia.org/wiki/File:Peixes_da_plan%C3%ADcie_de_inunda%C3%A7%C3%A3o_do_alto_rio_Paran%C3%A1_(3678425)_(Colossoma_macropomum).png), [Hoplosternum](https://commons.wikimedia.org/wiki/File:Atipa_from_French_Guiana.jpg), and [Hoplias](https://commons.wikimedia.org/wiki/File:Hoplias_malabaricus_(16521590825).jpg) used in Figure 5. We also thank the journals Papers in Palaeontology, Journal of Morphology, and Neotropical Ichthyology for allowing the reuse of pictures of the genera *Lepidosiren*, *Phractocephalus*, *Platysilurus*, *Zungaro*, *Hydrolycus*, *Piaractus*, *Hemidoras*, and *Oxydoras* (Ballen et al.; 2022, 2026), *Trachycorystes* (Britski & Akama, 2011), *Lophiosilurus* (Marques et al., 2025), and *Glanidium* (Sarmento-Soares & Martins-Pinheiro, 2013). We thank the editor Dr. Taissa Rodrigues, and two anonymous reviewers for their insightful comments.

## Notes

### Competing Interest Statement

The authors have declared no competing interest.

### Summary of Updates

We have updated the manuscript following suggestions during peer review. We are also adding a figure.

