## Supplementary material for "Acrean fish diversity back in time: Paleoichthyology of the Rio Acre fossil fauna in Brazil"

Gustavo A. Ballen[1,*], Karina Alencar[2], Fellipe Muniz[3], Edson Guilherme[4], Leonardo Aldrin[5], Silvia Lomba[2], Lucas R. Dias-Costa[2], and Annie S. Hsiou[2,*]

[1]Instituto de Biociências, Universidade Estadual Paulista "Júlio de Mesquita Filho", Câmpus Botucatu, Botucatu, SP, Brazil.

[2]Laboratório de Paleobiodiversidade Integrada, Departamento de Biologia, Faculdade de Filosofia, Ciências e Letras de Ribeirão Preto, Universidade de São Paulo, Ribeirão Preto, São Paulo

[3]Laboratório de Paleontologia de Ribeirão Preto, Departamento de Biologia, Faculdade de Filosofia, Ciências e Letras de Ribeirão Preto, Universidade de São Paulo, Ribeirão Preto, São Paulo

[4]Laboratório de Pesquisas Paleontológicas, Centro de Ciências Biológicas e da Natureza, Universidade Federal do Acre, campus Rio Branco, Acre.

[5]Laboratório de Macroevolução e Paleobiologia, Departamento de Zoologia do Instituto de Biociências da USP

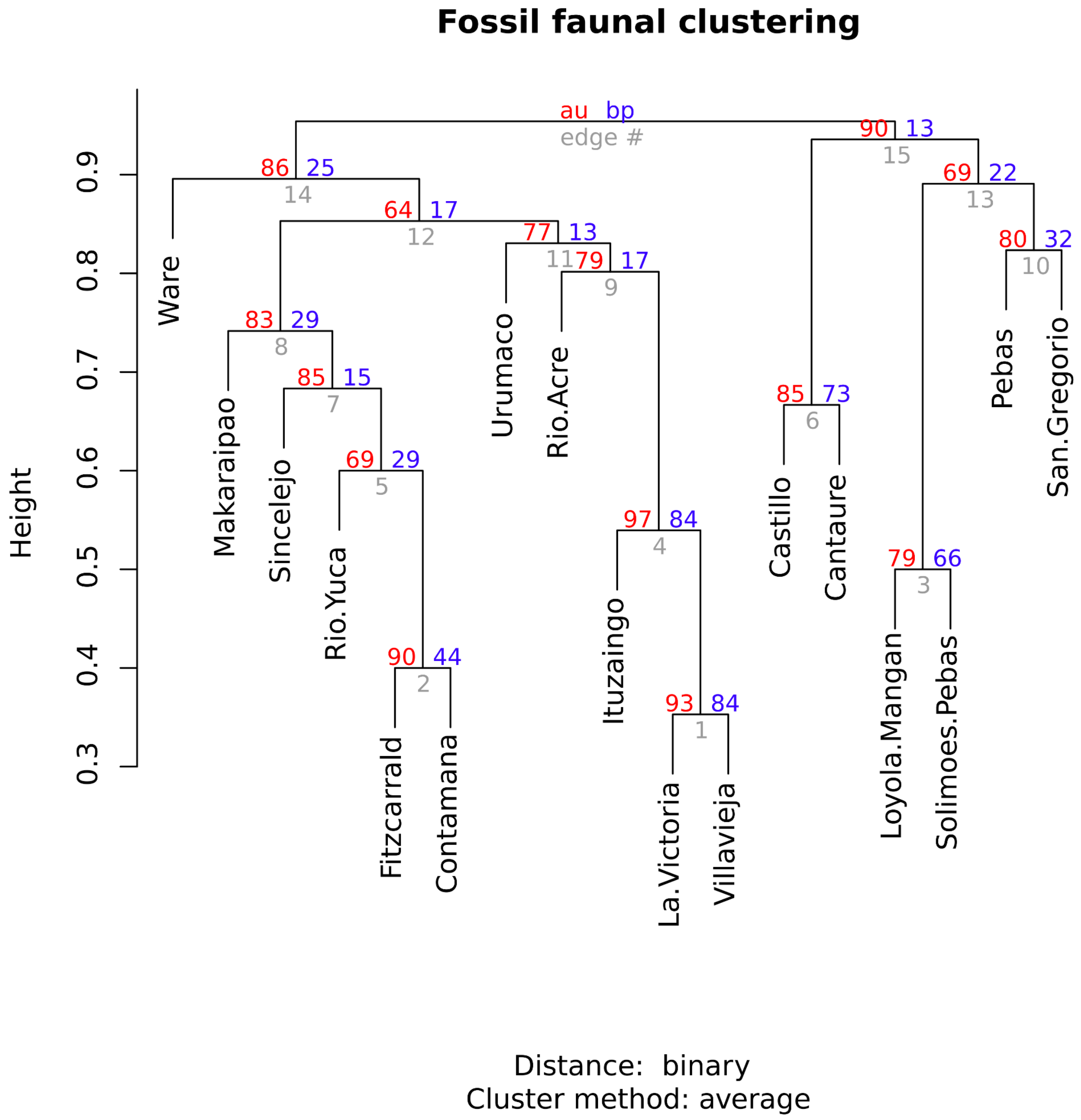


**Figure S1.** Faunal clustering of Neogene freshwater fish faunas in South America. Support values to the left and right of each node are approximately unbiased and bootstrap nodal support values, respectively.
